# Cryo-EM structure of S-Trimer, a subunit vaccine candidate for COVID-19

**DOI:** 10.1101/2020.09.21.306357

**Authors:** Jiahao Ma, Danmei Su, Xueqin Huang, Ying Liang, Yan Ma, Peng Liang, Sanduo Zheng

**Author notes:** These authors contributed equally (Jiahao Ma, Danmei Su).

## Abstract

Less than a year after its emergence, the severe acute respiratory syndrome coronavirus 2 (SARS-CoV-2) has infected over 22 million people worldwide with a death toll approaching 1 million. Vaccination remains the best hope to ultimately put this pandemic to an end. Here, using Trimer-Tag technology, we produced both wild-type (WT) and furin site mutant (MT) S-Trimers for COVID-19 vaccine studies. Cryo-EM structures of the WT and MT S-Trimers, determined at 3.2 Å and 2.6 Å respectively, revealed that both antigens adopt a tightly closed conformation and their structures are essentially identical to that of the previously solved full-length WT S protein in detergent. These results validate Trimer-Tag as a platform technology in production of metastable WT S-Trimer as a candidate for COVID-19 subunit vaccine.

---

The emergence of SARS-CoV-2 in late 2019 has led to a global pandemic and has disrupted lives and global economies on a scale unseen in recent human history. This is not the first time when a new coronavirus has posted as a major threat to public health; both SARS-CoV and Middle East Respiratory Syndrome (MERS-CoV) caused human infections within past 17 years (1). The fact that no licensed vaccines have ever been approved for these highly similar viruses is a reminder for the great challenges we face when hundreds of companies and institutions worldwide rush to develop COVID-19 vaccines with multiple strategies (2).

A successful vaccine that could truly impact the course of this ongoing COVID-19 pandemic has to have four key characteristics: safety, efficacy, scalability (to billions of doses to meet global demand), and speed. Although protein subunit vaccines have excellent track records for the first three requirements, exemplified by the highly successful vaccine Gardasil used to prevent HPV infections (3) and Shingrix vaccine for containing herpes zoster virus infections (4), subunit vaccine development can take years to decades to complete. Many of the difficulties reside in the manufacturing processes that have to ensure a fully native-like antigen structure is retained, starting from subunit vaccine designs. Similar to other enveloped RNA viruses such as HIV, RSV and influenza, coronaviruses including SARS-CoV-2 also use a ubiquitous trimeric viral surface antigen (Spike protein) to gain entry into host cells. In the case of SARS-CoV-2, this occurs via binding to ACE2 receptor expressed in target cells (5). Approaches for using a non-covalent trimer-foldon from bacterial phage GCN4 protein to stabilize the trimer conformation while simultaneously introducing mutations in viral antigens to abolish furin cleavage and stabilize the antigen in pre-fusion forms (*6, 7*) are common strategies inherited from decades of HIV and RSV vaccine studies, but these have yet to yield a successful vaccine.

Using Trimer-Tag technology (8), we produced both soluble wild-type (WT) and a furin site mutant (MT) (R685A) forms of S-Trimer from CHO cells in serum-free fed-batch processes in bioreactors with protein yield ranging from 0.5 −1 g/L (fig. S1A). These antigen titers are three orders of magnitude higher than that reported previously for foldon derived S proteins with two Pro mutations (S-2P) (6,7), laying a solid foundation for scalability requirement of a successful COVID-19 vaccine. Both S-Trimers consist of the ectodomain (amino acid residues 1-1211) of SARS-CoV-2 Spike protein fused in-frame to the C-terminal region of human type I (α) collagen, which spontaneously form a disulfide-bond linked homo-trimer, thereby stabilizing the antigens in trimeric forms (Fig.1A-B). Using a tailored affinity purification scheme that employs a collagen receptor Endo180-Fc fusion protein that binds to the Trimer-Tag with high affinity, the secreted S-Trimers were purified to near homogeneity in a single step (see accompanying paper). Reducing SDS-PAGE analysis of the purified S-Trimers revealed that the WT S-Trimer was metastable and partially cleaved precisely at S1/S2 boundary by furin, while a single point mutation (R685A) in MT S-Trimer fully abolished the protease cleavage (Fig.1C). WT spike proteins from live SARS-CoV-2 (9) or recombinant full-length S (10) were all previously shown to be partially cleaved apparently at S1/S2 boundary by furin proteases. In contrast, an S-Trimer derived from wild-type SARS-CoV-1 S-protein produced in the same manner was essentially uncleaved by furin protease, like the MT S-Trimer from SARS-CoV-2 (Fig.1C). Receptor binding studies using ForteBio BioLayer interferometry showed that all three S-Trimers had similar high affinity to ACE2-Fc with 1.2 nM (fig. S1B), similar to previous reports using purified S-2P protein (5). Furthermore, our finding of SARS-CoV-1 S-Trimer having a similar, if not higher affinity to ACE2-Fc than that of SARS-CoV-2, seems to be contradictory to the hypothesis that the difference in virulence between the two viral strains stems from difference in their receptor binding affinities (5). Instead, our finding supports that the much higher infectivity of SARS-CoV-2 is more likely attributed to furin cleavage of the spike protein that is largely absent in that of SARS-CoV-1 and other earlier stains of coronaviruses (11).

**Fig. 1.**
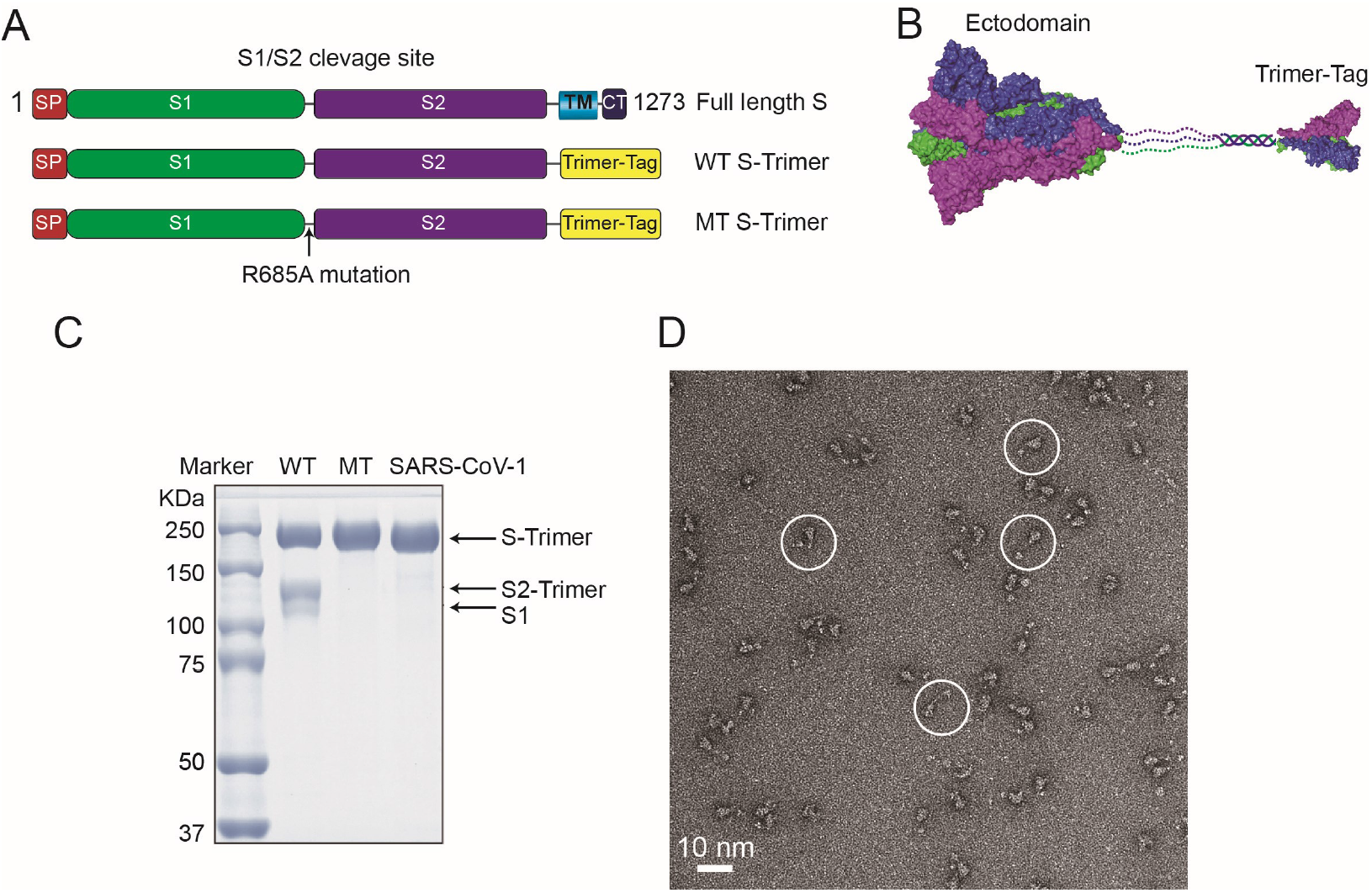
Structural design of the trimerized SARS-CoV-2 spike protein. (A) Schematic representation of the full-length spike protein, WT S-Trimer and MT S-Trimer. The ectodomain of full-length S is fused with a trimer tag derived from the C-terminal domain of human type I(α) collagen to produce WT S-Trimer. A single point mutation R685A at the S1/S2 cleavage site was introduced in the WT S-Trimer to generate MT S-Trimer. The full-length S protein includes: S1, S2, S1/S2 cleavage site, transmembrane domain (TM) and cytoplasmic tail (CT). (B) Cartoon representation of the WT and MT S-Trimer. Structure of the ectodomain of the spike protein (PDB: 6VXX) and the C-terminal domain of collagen (PDB ID: 5K31) are shown as surface. Dashed line represents region not resolved in the structure. (C) The purified WT S-Trimer, MT S-Trimer of SARS-CoV-2 and SARS-CoV-1 are analyzed by Coomassie stained SDS-PAGE. (D) A representative negative-stain EM image of the MT S-Trimer.

Negative staining electron microscopy (EM) analysis of the MT S-Trimer revealed homogeneous particles consistent with trimeric spike proteins connected to the Trimer-Tag (Fig. 1D). Particles with similar structural features have also been observed by negative-stain EM for WT S-Trimer (see accompanying paper).

To further investigate structural details of these vaccine candidates, we first sought to determine the cryo-EM structure of the MT S-Trimer which is more tractable for structural determination. While the particles showed preferred top-view orientation when embedded in ice, they preferentially adopted side view orientation on graphene oxide-coated grids (fig. S2). Combination of the two datasets enabled us to obtain a structure at 2.6 Å resolution (Fig. 2A and fig. S3). Due to the high resolution, the EM density for side chains was clear and water molecules could be observed in the structure (fig. S3D-3H and Table S1). In addition to all the glycosylation sites and disulfide bonds previously observed in the S-2P structure (PDBID: 6VXX), the disulfide bond between Cys15 and Cys136 and the N-linked glycan at Asn17, both in the NTD domains, were well resolved in our structure (fig. S4 and Table S2). The Trimer-Tag was invisible in our structure due to highly flexible nature of the linker between the soluble S and the C-prodomain of collagen. After 3D classification without imposing three-fold symmetry (C3), we found that all three RBD domains in our map adopt a closed conformation (RBD down) without any open conformation (RBD up) that was previously observed in the S-2P structure (6). Moreover, when the S2 domains were aligned, three S1 domains shift toward the three-fold axis compared to that in the S-2P structure, forming a tightly closed trimer (fig. S5). Surprisingly, we observed unaccounted for EM density in both the NTD and RBD region of the S1 domain. The presence of polysorbate 80 (PS80) during the purification process suggested that the bulky EM density in the NTD region could be accounted for by PS80 (fig. S3E). The EM density in the RBD region is elongated and in close proximity to R408, we speculate it may be oleic acid or linoleic acid which is present in the culture medium (fig. S3F). Indeed, PS80 and oleic acid could be well fitted into the density of the NTD domain and the RBD domain, respectively, owing to the excellent quality of EM map (Fig. 2A and fig. S3D-3F). Mass spectrometry analysis further confirmed their identities (fig. S6). Interestingly, instead of oleic acid seen in our structure, linoleic acid as well as PS80 were also observed in the recently published structure of a full-length mutant S protein (3Q-2P-FL) produced from insect cells (7). PS80 is buried deeply in the hydrophobic pocket residues with a few hydrophilic residues including N99, N121, R190 and H207 making hydrogen bonds with the hydroxyl group of the PS80 (Fig. 2B). Notably, PS80 engages hydrophobic interactions with F175 and M177 which are invisible in the S-2P structure (Fig. 2C). Since no small molecule was shown to bind to the S-2P structure, PS80 likely stabilizes the disordered loops of the NTD domain, making them more ordered (Fig. 2B and 2C). The oleic acid located in the hydrophobic pocket of the RBD domain engaged a salt bridge interaction with R408 at the adjacent protomer through its carboxylic acid group, bringing the RBD domain in close proximity and resulting in the tightly closed conformation (Fig. 2D).

**Fig. 2.**
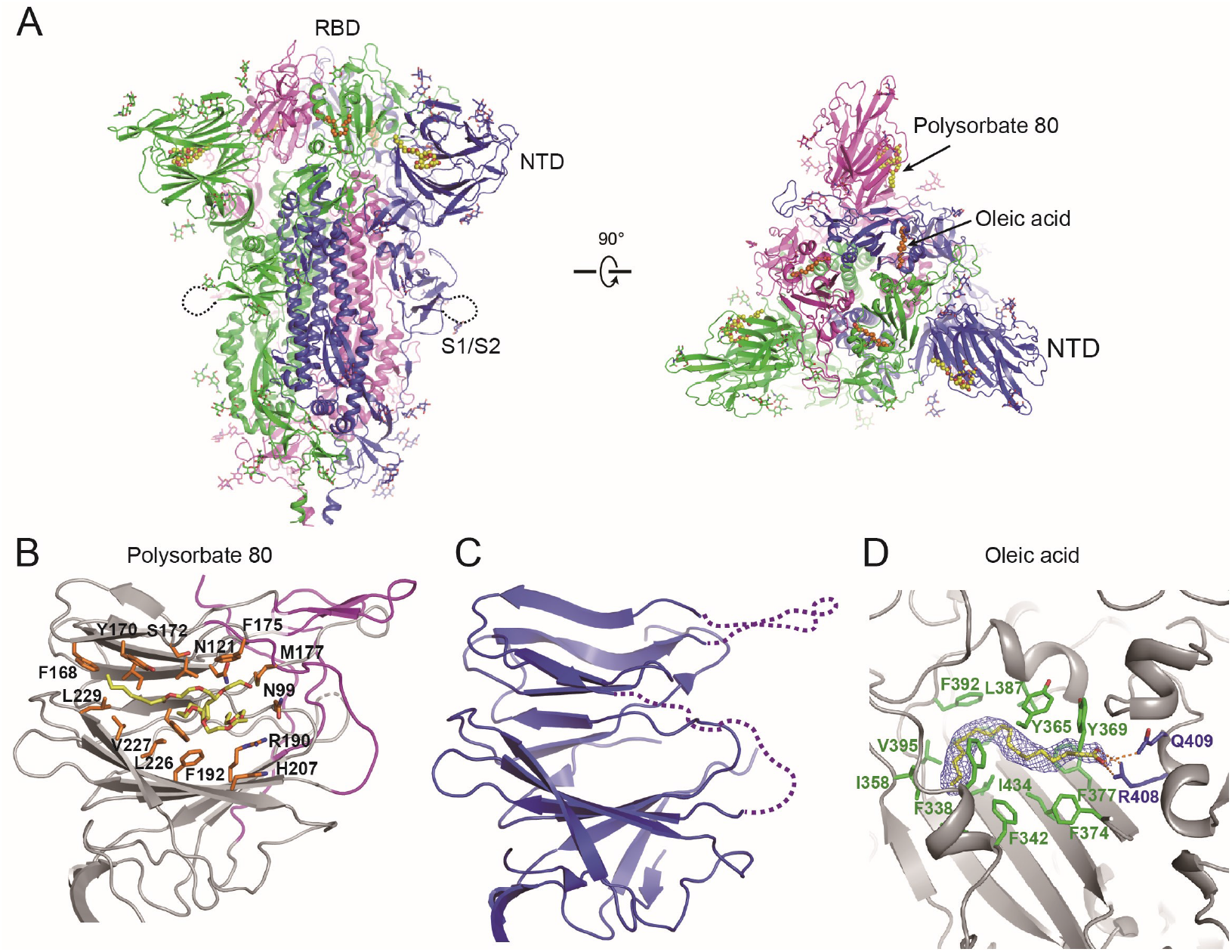
Cryo-EM structure of the MT S-Trimer. (A) Ribbon representation of the MT structure colored by subunit from two orthogonal views. Oleic acid and PS80 are shown as spheres and colored in orange and yellow respectively. (B) Detailed view of the NTD domain bound to the PS80. Structure colored by magenta corresponds to the flexible region at the NTD of the S-2P structure. (C) Structure of the NTD domain of the S-2P (PDB ID: 6VXX). Dashed line represents disordered region. (D) Oleic acid bridges the two adjacent RBD domains. Oleic acid is colored in yellow and the side chains of the two adjacent RBDs colored in green and blue respectively. The EM density for oleic acid is shown.

Recent studies have shown that low pH can stabilize the S-Trimer (12). Indeed, negative staining EM analysis of WT S-Trimer at pH 5.5 revealed more homogenous trimer than that at physiological pH (See accompanying paper). In light of this finding, we were able to determine the cryo-EM structure of the WT S-Trimer at 3.2 Å resolution at pH 5.5 (Fig. S7 and Table S1). The structure of the WT S-Trimer resembled that of the MT form with a root mean square deviation of 0.5 Å over 2773 Cα atoms (Fig. 3A). Oleic acid was well resolved in the WT structure but the density for the PS80 was weak, likely due to the low resolution or low occupancy. As a result, the NTD domain of the WT S-Trimer was less well resolved than that of MT (Fig. 3A). It has been shown that a pH-dependent switch domain (residue 824-848, pH switch 1) undergoes dramatic conformational change at different pH values (12). However, this region was nearly identical between our two structures (Fig. 3B). Instead, a fragment (residue 617-639) we named pH switch 2 at the CTD1 region of the S1 domain before the furin cleavage site displays significant structural arrangement. Whereas this region appeared disordered in the MT structure at physiological pH conditions, it was well ordered and formed a helix-turn-helix structural motif in the WT structure at pH 5.5 (Fig. 3C). From structural perspective, lower pH likely contributes to this structural arrangement. At physiological pH, R319 forms salt bridge interactions with D737 and D745 (Fig. 3D). At lower pH, the protonation of D737 and D745 weakens these interactions. As a result, R319 flips to the other side and makes hydrophobic interactions with W633 and L629 through its aliphatic chain, leading to the ordered helix-turn-helix motif (Fig. 3E). The newly-formed structural motif makes direct contact with the previously identified pH switch 1 of the adjacent protomer, accounting for the enhanced stability of the WT S-Trimer at lower pH (Fig. 3A). The structural arrangement of pH switch 2 in different pH was also observed in previous studies (12), further supporting the conformational change between the MT and WT S-Trimer structures was due to the different pH but not to the mutation in the furin site.

**Fig. 3.**
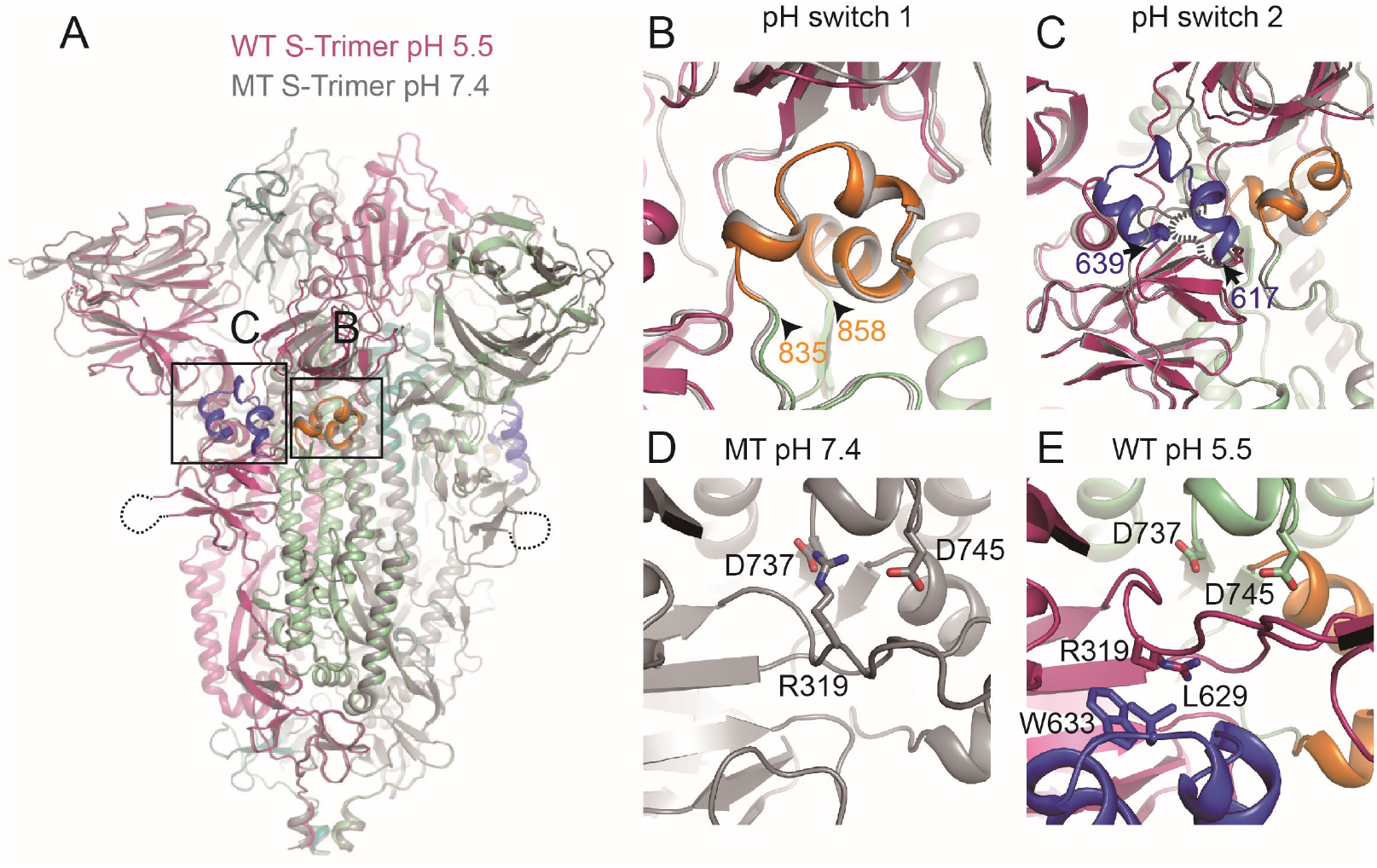
Cryo-EM structure of the WT S-Trimer at pH 5.5. (A) Structural overlay of the MT and WT S-Trimer. MT was colored in grey, and WT colored by subunits. Two pH switches in the WT S-Trimer structure are boxed and colored in blue and orange respectively. Comparison of pH switch 1 (B) and pH switch 2 (C) between MT and WT S-Trimer. Dashed line represents disorder region in pH switch 2 of the MT structure. (D) pH switch 2 is flexible in the MT structure where R319 makes electrostatic interactions with D737 and D745. (E) At low pH, the switch 2 forms a helix-turn-helix motif in the WT structure in which R319 flips and stabilizes it.

In contrast to the structural differences described above for S-2P protein, both of our WT and MT S-Trimer were nearly identical to the recently published structures of full-length wild-type S (10) and 3Q-2P-FL (7) purified in detergent from HEK293 and sf9 insect cell membranes, respectively. When revisiting the electron density map for full-length wild-type S protein (EMDB: 22292), we spotted unassigned density at the same position as oleic acid which was absent in the S-2P map (EMDB: 21452) (fig. S8). Therefore, fatty acid binding at the RBD domain may stabilize the tightly closed conformation, accounting for the conformational difference from S-2P. Based on the structural similarity among all published structures of SARS-CoV-2 S protein, we can classify them into three distinct states: a tightly closed state, a loosely closed state and an open state (Fig. 4A). A conformational switch (residue 835-858) previously known as the pH switch or the fusion-peptide proximal region (FPPR) is critical for conformational transition from the tightly closed state to the open state (Fig. 4B). In the tightly closed state which is less accessible to the receptor, three RBDs are down and in closer proximity to one another. The conformational switch region is well ordered and stabilizes this tightly closed state. Recently, D614G mutation has become predominant over the ancestral form worldwide and has been shown to increase viral infections (*13, 14*). In the tightly closed state, D614 makes a salt bridge interaction with K854 at the conformational switch region (Fig. 4C). From the tightly to the loosely closed state, the conformational switch undergoes a large conformational arrangement and becomes disordered (Fig. 4D). K854 flips to the opposite side and interacts with D568 and D574 of the CTD1, causing the S1 to move downwards relative to the S2 (Fig. 4A). Finally, the CTD1 domain further moves downwards and causes the RBD to adopt an open conformation for receptor binding (Movie S1). Therefore, D614G mutation abolishes the salt bridge interaction with K854 which in turn engages the CTD1 domain, thus favoring the open state.

**Fig. 4.**
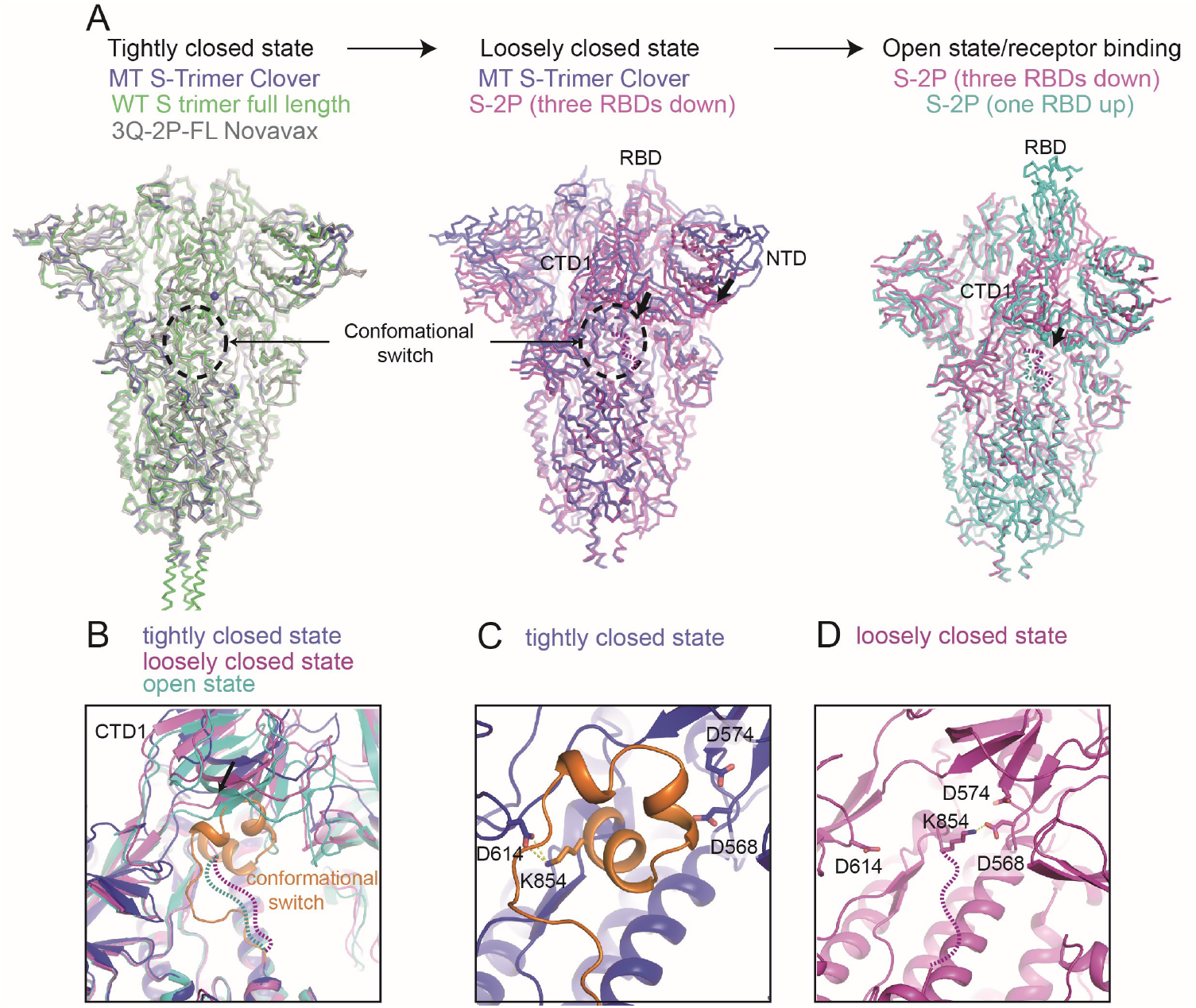
Conformational change of the SARS-CoV-2 spike protein before engaging ACE2 receptor. (A) Conformation transition from the tightly closed conformation (WT/MT structure) to the loosely closed conformation (PDB ID: 6VXX), and finally to the open conformation (PDB ID: 6VYB). The direction of movement of the NTD and CTD1 of the S1 domain during conformational change is marked with an arrow. The conformation switch region boxed is disorder in both the loosely closed and open conformation and is represented as dashed line. (B) A close-up view of the conformational switch in three conformations. The ordered switch would clash with CTD1 domain in the loosely closed and open conformation (C) K854 in the switch region engages D614 in the tightly closed conformation. (D) In the loosely closed conformation, the switch region undergoes structural arrangement, in which K854 flips and pulls S1 domain downwards through interacting with D574 and D568 in the CTD1 region.

To our knowledge, this is the first cryo-EM structure of the wild-type S protein in soluble and cleavable form without the transmembrane domain, confirming structural integrity of this metastable wild-type form of COVID-19 subunit vaccine candidate. While most vaccine candidates currently in clinical trials incorportated furin site and double proline mutations, our wild-type S-Trimer offers a more native-like antigen for potentially more effective and broader protection against SARS-CoV-2 infection. Thus, Trimer-Tag technology that has been proven here to be able to rapidly produce large quantities of native-like S-Trimer antigen, may offer a platform technology for subunit vaccine development for enveloped RNA viruses that use ubiquitous trimeric antigens to invade host cells.

Like the previously reported structure of full-length WT S protein purified in detergent micelles, it is unclear whether the furin cleavage site in our resolved WT S-Trimer structure is cleaved. Moreover, we could not exclude the possibility that other conformational states exist in the WT sample that were not captured in our cryo-EM study since partial cleavage of the furin site may lead to some S1 dissociation from S-Trimer. Nevertheless, we are certain that the highly purified WT S-Trimer predominately adopts a pre-fusion state, unlike the full-length wild-type spike protein which forms both pre- and post-fusion states in the presence of detergent (10).

Consistent with structural studies reported here showing that WT S-Trimer is native-like, preclinical studies showed that this COVID-19 vaccine candidate resulted in rapid and high-level induction of both neutralizing antibodies and Th1-biased cellular immune responses in mice, rats and nonhuman primates. Non-human primates immunized with WT S-Trimer were fully protected from SARS-CoV-2 viral challenge (See accompanying paper). Currently, an S-Trimer subunit vaccine candidate is under clinical investigation.

## ACKNOWLEGEMENTS

We thank Xiaodong Wang for his coordination and input in this study. We thank Maofu Liao and Andrew C. Kruse for critical reading of the manuscript. We thank Hongwei Wang for providing graphene oxide coated grids. We also thank staff at Shuimu BioSciences for their assistance with cryo-EM data collection. All EM data were collected at Shuimu BioSciences.

## Funding

This work was supported by grants from Coalition for Epidemic Preparedness Innovations (CEPI), Chinese Ministry of Science and Technology, Beijing Municipal Commission of Science and Technology, Tsinghua University and Chengdu Bureau of Science & Technology (2020-YF08-00024-GX).

## Author Contribution

D. S. expressed and purified the S-Trimers, J.M. collected negative stain EM images. J.M and S.Z. prepared cryo grids and collected cryo-EM data. X. H and Y. L performed ForteBio affinity analysis. S.Z. performed cryo-EM data processing and model building. Y.M. did mass spectrometry experiments. S.Z. and P.L conceived this project, directed the experiments and wrote the manuscript with input from all other authors.

## Competing interests

P.L. has ownership interest in Clover Biopharmaceuticals. All other authors declare no competing interests.

## Data and materials availability

The atomic structures have been deposited at the Protein Data Bank (PDB) under the accession codes XXX and XXX. The EM maps have been deposited at the Electron Microscopy Data Bank (EMDB) under the accession numbers XXX and XXX.

## Supplementary Materials for

### This PDF file includes

Materials and Methods

Figs. S1 to S8

Tables S1 to S2

Movies S1

## Materials and Methods

### Protein expression and purification

Endo180-Fc expression vector was generated by subcloning a PCR amplified cDNA encoding soluble human Endo180 (amino acid residue 1-1394) (Thomas et al) into the *HindIII* site of pGH-hFc expression vector (GenHunter, Nashville, TN) to allow in-frame fusion to human IgG Fc. The expression vector was transfected into GH-CHO *(dhfr-/-)* cell line (GenHunter, Nashville, TN) using FUGENE 6 (Roche) and grown in IMDM medium with 10% FBS. After stepwise gene amplification with increasing concentrations (0.0–10 nM) of MTX (Sigma), a high titer clone was then adapted to SFM-4CHO serum-free medium (GE BioSciences). The secreted End0180-Fc fusion protein was then produced in a 15 L bioreactor (Applikon) under a fed-batch process with Cell Boost 2 supplement (GE Hyclone) as instructed by the manufacturer. Endo180-Fc was purified to homogeneity by protein A affinity chromatography using MabSelect PrismA (GE Healthcare) followed by Capto QXP resins (GE BioSceinces) in a flowthrough mode to remove any host cell DNA and residual host cell proteins (HCP).

To produce S-Trimer fusion proteins, cDNA encoding the ectodomain of either wild-type SARS-CoV-2 S protein (amino acid residues 1 to 1211) (GenBank: MN908947.3) or a SARS-CoV (amino acid residues 1 to 1193) (GenBank: AAS00003.1) were synthesized using -optimized codons for *Cricetulus griseus* (CHO Cell) by GenScript. The cDNAs were subcloned into pTRIMER expression vector (GenHunter, Nashville, TN) at *Hind III* and *Bgl II* sites to allow in-frame fusion of the soluble S protein to Trimer-Tag as described previously (1). Furin cleavage site mutant S-Trimer (R685A) was generated by site-directed mutagenesis using mutagenesis primer pair (5’-gcccaaggagggctgcgtctgtggctagcc-3’ and 5’-ggctagccacagacgcagccctccttgggc-3’) and WT S-Trimer expression vector as a template following the protocol of QuikChange kit (Strategene). The expression vectors were transfected into GH-CHO *(dhfr-/-)* cell line (GenHunter, Nashville, TN) using FUGENE 6 (Roche) and grown in IMDM medium with 10% FBS. After stepwise gene amplification with increasing concentrations (0.0–10 nM) of MTX (Sigma), clones producing the highest S-Trimer titer were adapted to SFM-4CHO serum-free medium (GE BioSciences). The secreted S-Trimer fusion proteins were produced in a 15 L bioreactor (Applikon) under a fed-batch process with Cell Boost 2 supplement (GE Hyclone) as instructed by the manufacturer.

Cell culture medium was clarified by depth filtration (Millipore) to remove cell and debris. S-Trimers were purified to homogeneity by consecutive chromatographic steps including Protein A affinity column using MabSelect PrismA (GE Healthcare) which was preloaded with Endo180-Fc at (3 mg/mL) to capture S-Trimer, based on the high affinity binding between Endo180 and Trimer-Tag (2). After washing off any unbound contaminating proteins, S-Trimers were purified to near homogeneity in a single step using 0.5 M NaCl in phosphate buffered saline (PBS). For S MT and SARS-CoV S-Trimer, the proteins were dialyzed against PBS plus 0.02% Polysorbate 80 before analysis. After one hour of low pH (pH 3.5) viral inactivation (VI) step using acetic acid, the pH was adjusted to neutral range, WT S-Trimer was further purified on a Capto QXP resins (GE BioSceinces) in a flow-through mode to remove any host cell DNA and residual host cell proteins (HCP). A final preventative viral removal (VR) step was performed using a nano-filtration cartridge (AsahiKASEI) before final buffer exchange to PBS plus 0.02% Polysorbate 80 by UF/DF (Millipore).

ACE2-Fc expression vector was generated by subcloning a gene-synthesized cDNA template (GenScript) encoding soluble human ACE2 (amino acid residue 1-738, accession number: NM_001371415.1) into *Hind III* and *Bgl II* sites of pGH-hFc expression vector (GenHunter, Nashville, TN) to allow in-frame fusion to human IgG Fc. The expression vector was then stably transfected into GH-CHO (dhfr -/-) cell line and high expression clones were selected and adapted to SFM-4-CHO (Hyclone) serum free medium and ACE2-Fc was produced in a 15 L bioreactor as essentially as described for S-Trimer above. ACE2-Fc was purified to homogeneity from the conditioned medium using PoRos XQ column (Thermo Fisher) following manufacturer’s instructions.

### Receptor binding studies of S-Trimers to human ACE2-Fc

The avidity of different S-Trimer binding to the SARS-CoV-2 receptor ACE2 were assessed by Bio-Layer Interferometry measurements on ForteBio Octet QKe (Pall, New York). ACE2-Fc (10 μg/mL) was immobilized on Protein A (ProA) biosensors (Pall). Real-time receptor binding curves were obtained by applying the sensor in a two-fold serial dilutions of S-Trimer from 22.5-36 μg/mL in PBS. Kinetic parameters (K_on_ and K_off_) and affinities (K_D_) were analyzed using Octet software, version 12.0. Dissociation constants (K_D_) were determined using steady state analysis, assuming a 1:1 binding model for a S-Trimer to ACE2-Fc.

### Negative staining

Negative staining was performed as previously described (3). In brief, 3 μl of purified S-trimer at a concentration of about 0.01 mg/ml was deposited on a glow-discharged carbon-coated copper grid for 30 s before being blotted with filter paper. Grids were quickly washed with two drops of water and one drop of 2% (w/v) uranium acetate. Grids were kept touching to the last drop of 2% (w/v) uranium acetate for 90 s, and blotted with filter paper. Data collection was performed on a Tecnai T12 electron microscope operated at 120 Kev equipped with a FEI Ceta 4K detector. Images were collected at a magnification of 57,000 x and a defocus of 1.5 μm.

### Cryo-EM sample preparation and data collection

Purified MT S-trimer protein diluted to 0.2 and 0.5 mg/ml in PBS buffer were applied to glow-discharged gold holey carbon 1.2/1.3 300-mesh grids with and without graphene oxide, respectively. Grids were blotted for 2-4 seconds at a blotting force of 4 and plunge-frozen in liquid ethane using a MarkIV Vitrobot (Thermo Fisher Scientific). The chamber was maintained at 8 °C and 100% humidity during freezing. 0.3 mg/ml of WT S-trimer sample in low pH buffer (100 mM sodium citrate pH 5.5 and 100 mM NaCl) was deposited on glow-discharged gold holey carbon 1.2/1.3 300-mesh grids with graphene oxide. Grids were blotted and vitrified using the same condition.

All movies were collected using a Titan Krios microscope (Thermo fisher Scientific) equipped with a BioQuantum GIF/K3 direct electron detector (Gatan). The detector is operated in superresolution mode. A complete description of cryo-EM data collection parameters are summarized in Table S1.

### Cryo-EM Image processing

For MT S-Trimer protein, motion correction for cryo-EM images and contrast transfer function (CTF) estimation were performed using motioncorr2 *(4)* and CTFFIND4 (5) respectively. 574,832 particles were automatically picked from 534 images collected on grid without graphene oxide (GO) using Laplacian-of-Gaussian in Relion 3.0.7 (6), and 1,029,938 particles was automatically picked from 1584 images collected on grid with GO. Extract particles from two datasets were downsized by two-fold and subjected to 2D classification separately, resulting in 438,205 and 560,545 good particles. Particles from GO grids were recentered using scripts written by Kai Zhang (http://www.mrc-lmb.cam.ac.uk/kzhang/useful_tools/scripts/) before 3D classification. Good particles from both datasets were combined and subjected to 3D classification using the initial model generated from the model of S protein with C1 symmetry (PDB ID: 6vxx). Two major classes accounting for 26.7% and 25.8% particles show clear and complete structural features. These particles were auto-refined followed by local 3D classification with C1 symmetry to generate two classes with similar structure. The better class was subjected to 3D refinement with C3 symmetry and post-processed using mask on the entire molecule to yield a 2.9 Å map. CTF refinement was performed to further increase the resolution to 2.6 Å.

For image processing of WT S-Trimer, MT S-Trimer map was low-pass-filtered to 20 Å resolution and used as the 3D reference template for auto-picking. 752,204 articles were picked from 1199 images collected on GO-coated grid and used for two rounds of 2D classification. 541,528 particles were selected from good 2D classes and used for 3D classification with C1 symmetry. One class accounting for 32.2% showing a well-defined structure was refined with C3 symmetry and post-processed using mask on the entire molecule to give a map at 3.4 Å resolution. CTF refinement followed by another round of 3D refinement improved the resolution to 3.2 Å. Reported resolutions were calculated based on the gold-standard Fourier shell correlation (FSC) at the 0.143 criterion.

### Model building

The soluble ectodomain structure S-2P (PDB: 6VXX) was used as a template for model building. The missing region in the S-2P structure can be de novo modeled in the MT S-Trimer map, owing to its high resolution. The model was manually built in COOT (7) and real space refinement was performed in Phenix (8) using rotomer, Ramachandran and secondary structure restraints. The model for WT S-trimer was built in COOT based on the MT structure and refined in Phenix using the same strategy. Refinement statistics are summarized in Tabel S1.

### Small molecules extraction from protein samples

20 μL of WT and MT S-Trimer samples at 1.21 mg/ml and 1.02 mg/ml respectively were transferred to Eppendorf tubes and placed in a heater block at 100 *°C* for 5 min. Samples were then extracted with 100 μL methanol. The tubes were vortexed and centrifuged at 15,000 rpm at 4 °C for 10 min. The supernatant was collected, transferred to glass vials, and 1 μL was injected for the LC-MS analysis. For oleic acid and linoleic acid reference samples, 10 mM stock solution of compounds were prepared in DMSO and diluted to 1 μM by methanol, and 1 μL was injected for the LC-MS analysis.

### LC-MS Analysis

LC-MS analysis of PS 80 and MT S-Trimer sample were performed by an Agilent 1290 UHPLC coupled to an Agilent quadrupole-time of flight (QTOF) mass spectrometer via an electrospray ionization source (ESI) with JetStream technology. The separation was performed by flow injection using isocratic flow of a solvent composed of 0.1% formic acid in 60% acetonitrile and 40% water. The flow rate was set at 0.2 mL/min for 0.5 min. Mass spectra were recorded in the positive ionization mode over a mass range from *m/z* 100 to 1500. The scan parameters were capillary voltage 4.0 kV and fragmentor 135V. The nitrogen pressure and flow rate on the nebulizer were 40 psi and 5 L/min, respectively. Other ion source parameters included drying gas temperature of 325 °C, sheath gas temperature of 350 °C and sheath gas flow rate of 10 L/min.

The relative quantitation for oleic acid and linoleic acid was performed by a Thermo Vanquish UHPLC coupled to a Thermo Q Exactive HF-X hybrid quadrupole-Orbitrap mass spectrometer. The chromatographic separation was performed using a Waters CSH C18 column (2.1×100 mm, 1.7 μm) and maintained at 40°C. The separation was performed using isocratic flow of a solvent composed of 90% acetonitrile, 10% water, and 2 mM ammonium acetate. The flow rate was set at 0.2 mL/min for 10 min. Full-scan-ddMS^2^ mass spectra were acquired in the range of 100-1500 *m/z* with the following ESI source settings: spray voltage 2.5 kV, aux gas heater temperature 380 °C, capillary temperature 320 °C, sheath gas flow rate 30 unit, aux gas flow gas 10 unit in the negative mode. MS1 scan parameters included resolution 60000, AGC target 3e6, and maximum injection time 200 ms while dd-MS2 parameters included resolution 30000, ACG target 2e5, maximum injection time 100 ms, isolation window 4.0 m/z, NCE 30.

---

**Fig. S1.**
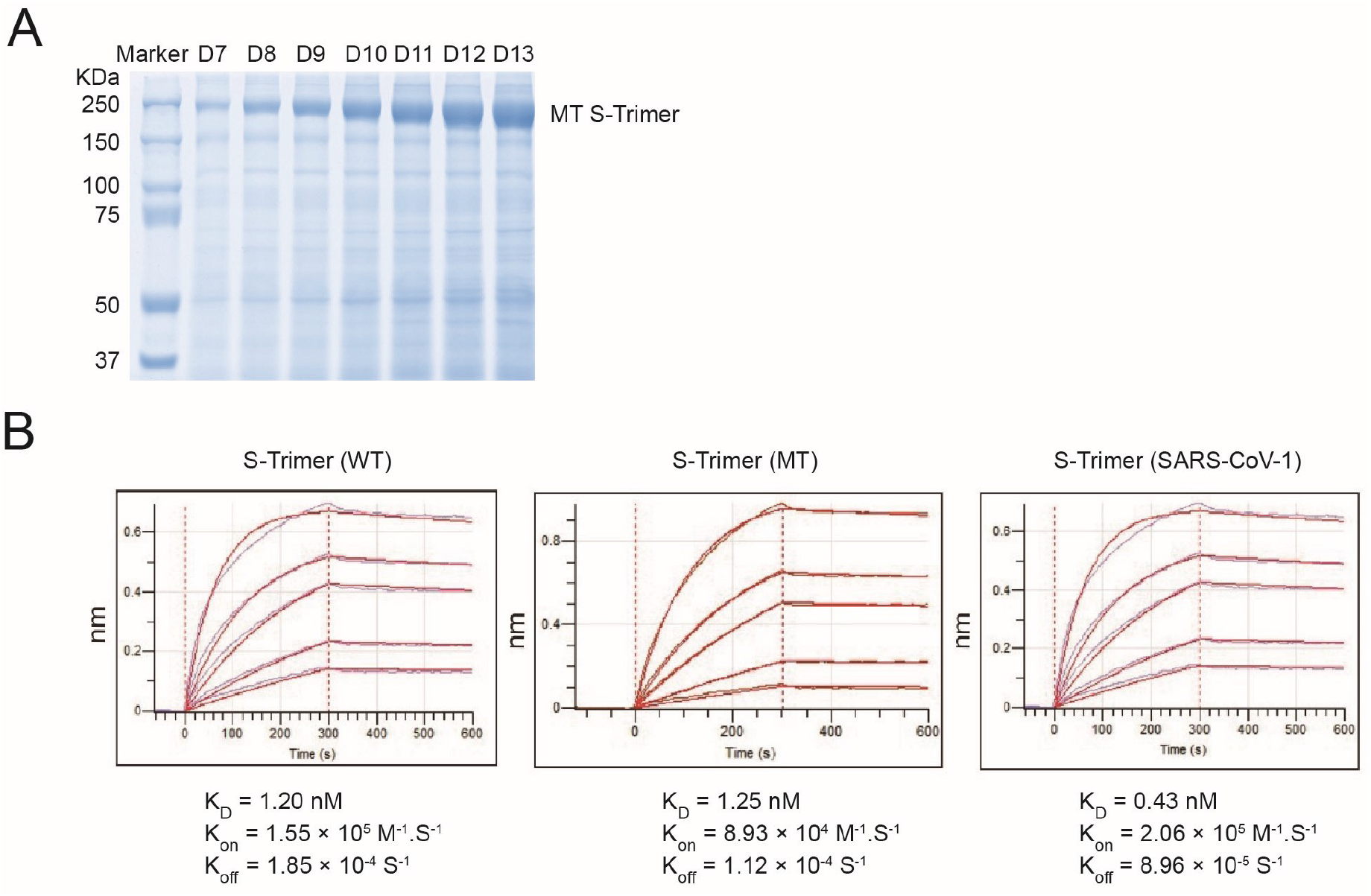
Measure the binding affinity between S-Trimer and Fc-tagged hACE2 ectodomain by ForteBio Bio-Layer Interferometry. (A) 10 μl of cell culture medium for MT S-Trimer expression was collected from day 7 (D7) to day 13 (D13) and separated by SDS-PAGE followed by Coomassie blue staining. (B) Receptor binding for WT and MT S-Trimer of SARS-CoV-2 and wild-type SARS-CoV-1 S-Trimer were analyzed as indicated.

**Fig. S2.**
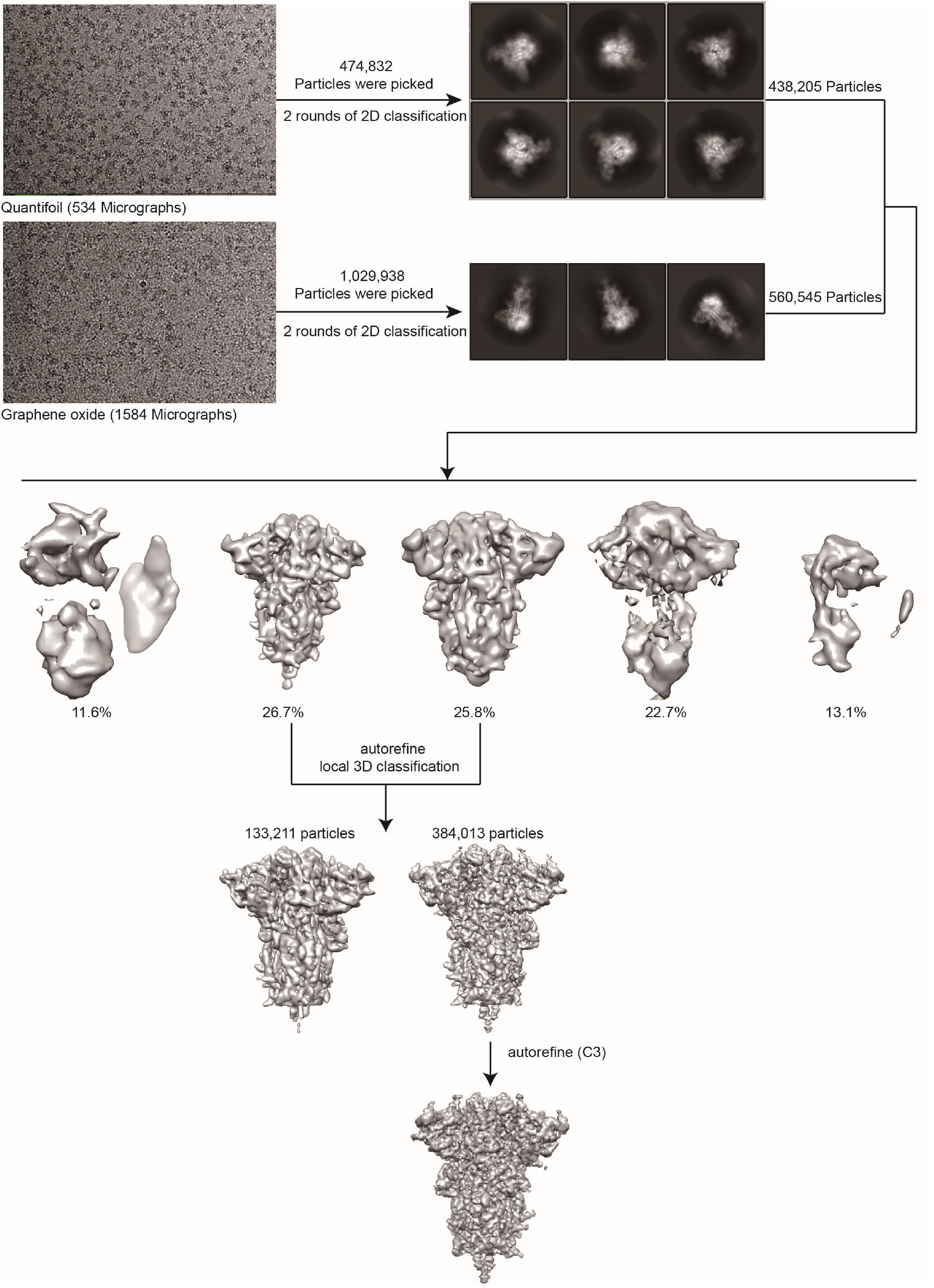
Cryo-EM data processing workflow for the MT S-Trimer.

**Fig. S3.**
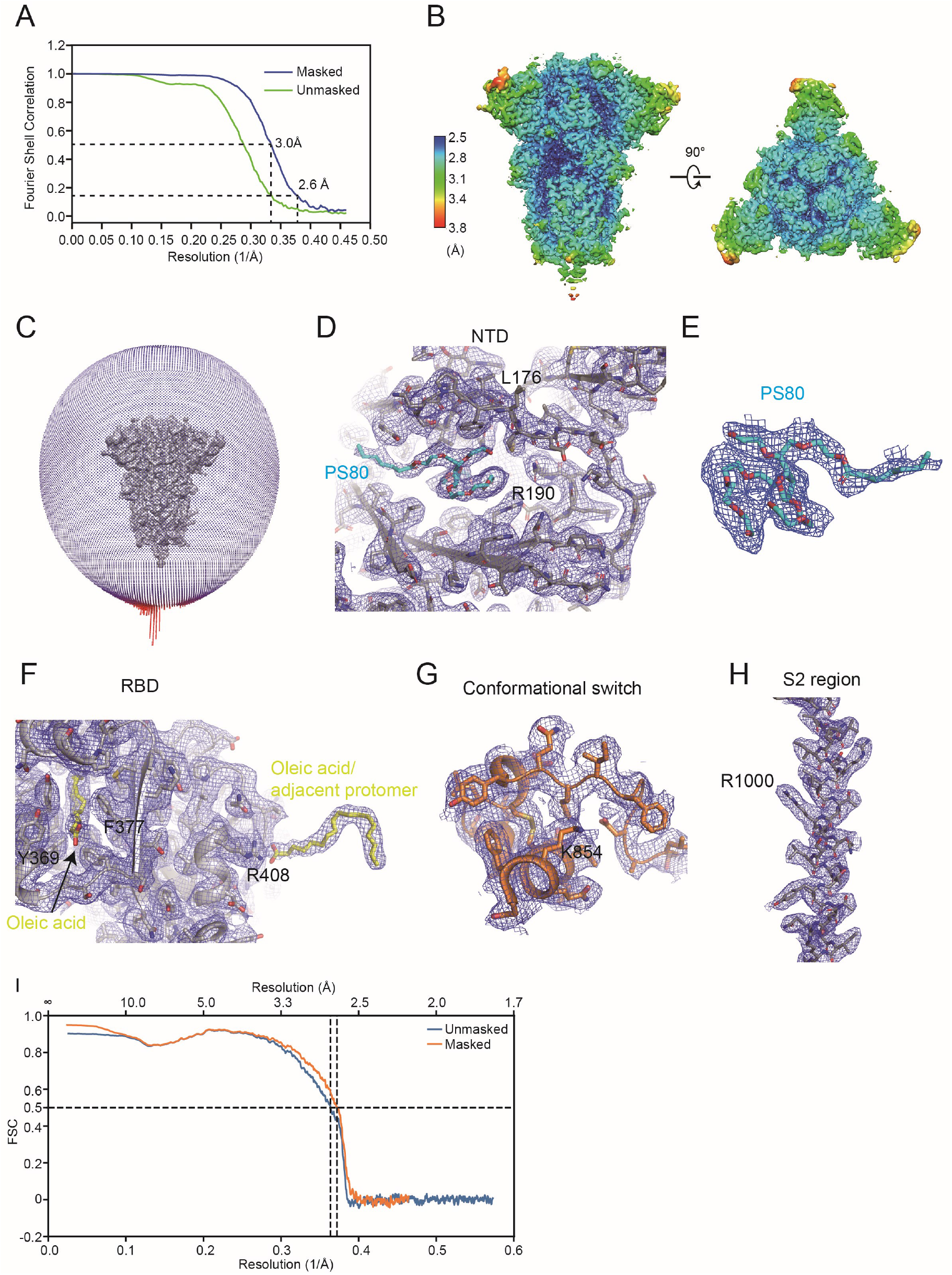
Cryo-EM structure validation for MT S-Trimer. (A) Gold standard FSC curves for the MT S-Trimer. Resolutions at FSC value of 0.5 and 0.143 were indicated on the masked FSC curve. (C) Angular distribution of the particles used for 3D reconstruction. Representative B-factor sharpened EM density map for the NTD (D), polysorbate 80 (E), the RBD with oleic acid (F), the conformational switch (G) and the S2 region (H). (I) FSC curves of model-to-map.

**Fig. S4.**
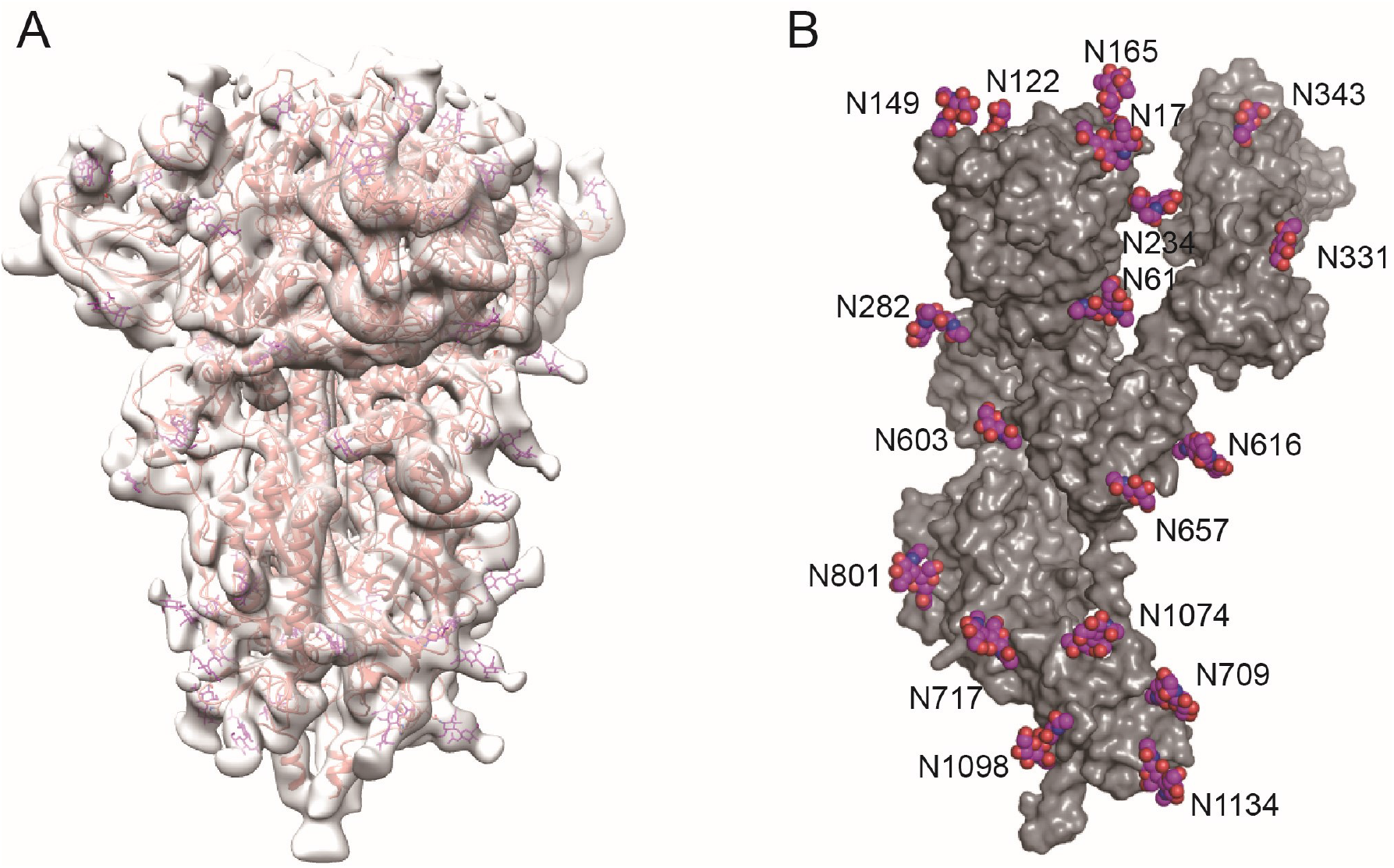
N-linked glycosylation sites on the MT S-Trimer structure. (A) Superposition of the low resolution EM density map and the MT S-Trimer model. The model is shown in ribbon and glycans are shown as magenta sticks. (B) Glycans are shown as sphere on the surface of one protomer and labeled with residue numbers of the linked asparagine.

**Fig. S5.**
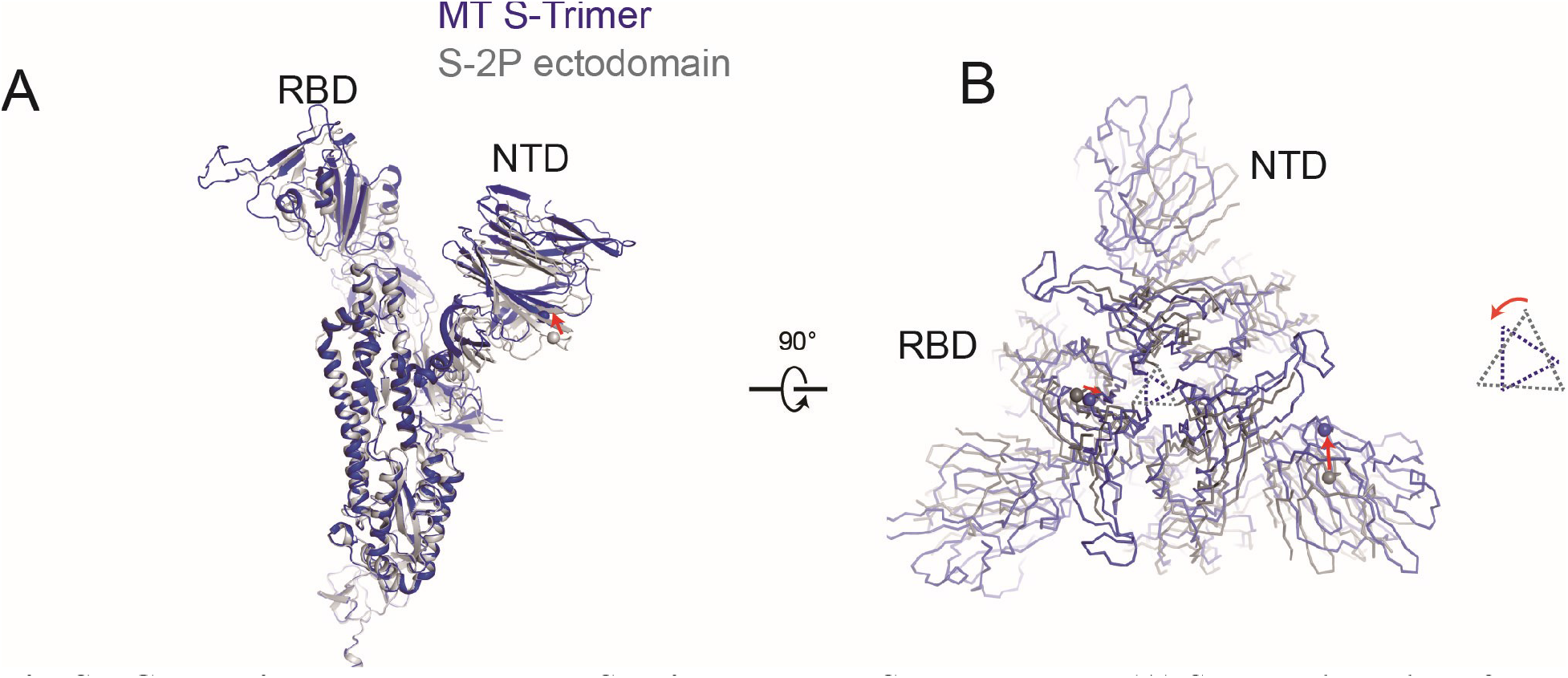
Comparison between the MT S-Trimer and the S-2P structure. (A) Structural overlay of MT S-Trimer and S-2P (PDB ID: 6VXX) with the S2 domain aligned. S-2P and MT are colored in grey and blue respectively. The same atoms from both structures are shown as sphere. The red arrow indicates moving direction. (B) Top view of the aligned MT and S-2P trimer structure. The RBD domains of the S-2P structure rotate counterclockwise and move toward three-fold axis relative to that of the MT structure.

**Fig. S6.**
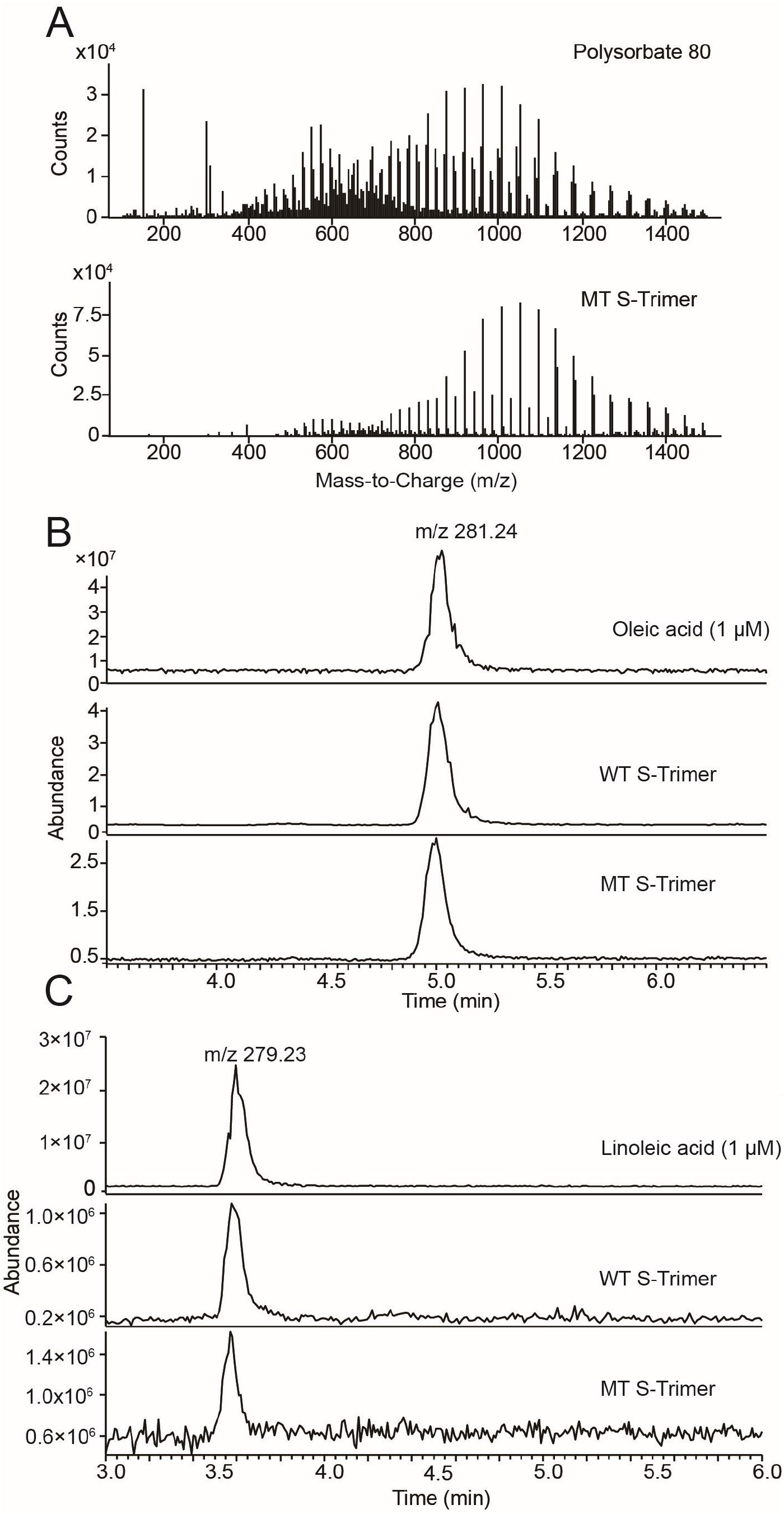
Small molecule identification of the purified S-Trimer sample by mass spectrometry. (A) Mass spectra of polysorbate 80 and small molecules extracted from the MT S-Trimer sample revealed that they are similar. (B) Extracted ion chromatograms (EICs) for small molecules from both the WT and MT S-Trimer with 1 μM of oleic acid (B) and linoleic acid (C) as references showed that S-Trimers were enriched with oleic acid.

**Fig. S7.**
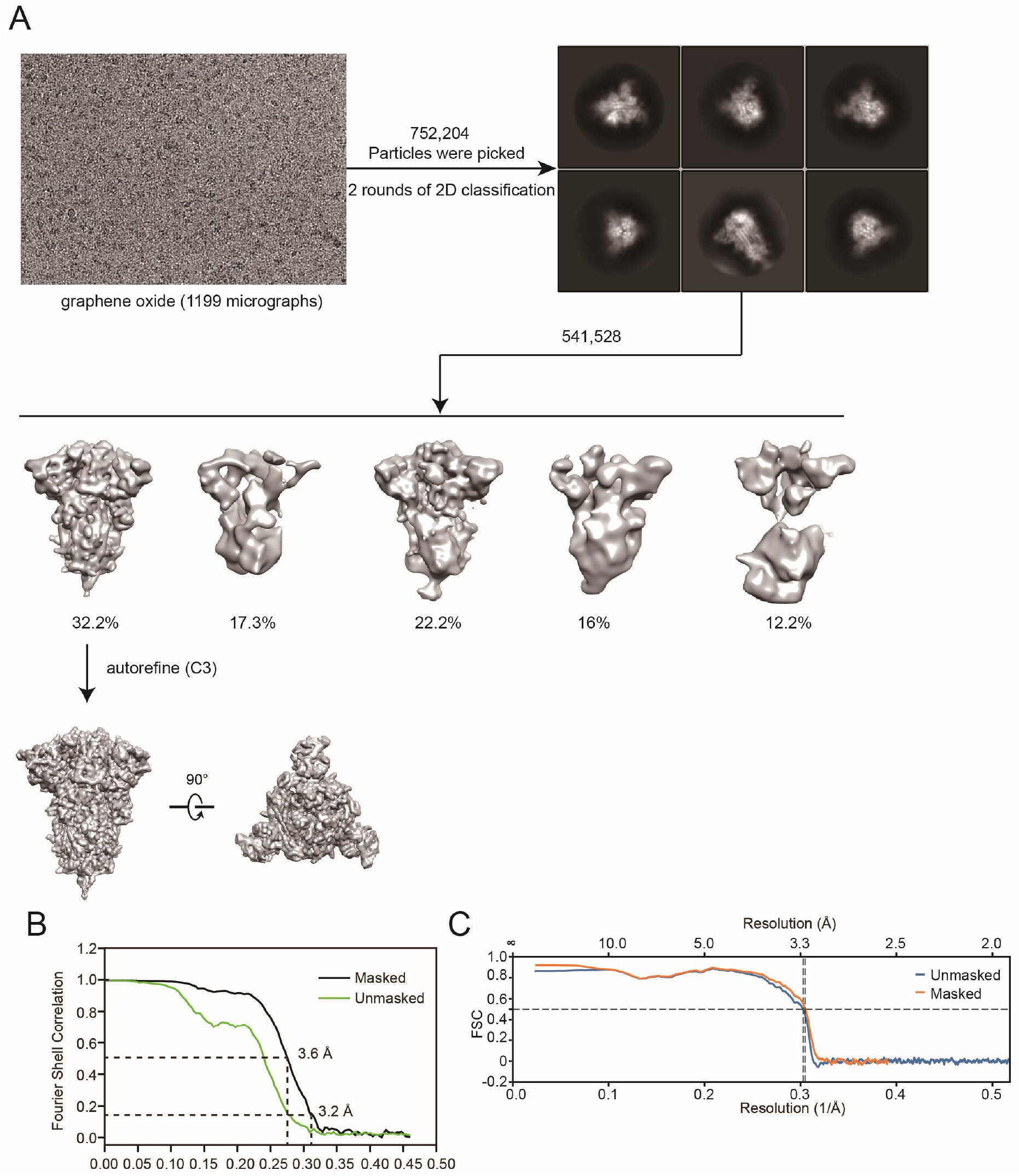
Cryo-EM data processing workflow for the WT S-Trimer. (A) Representative micrograph on graphene oxide, selected 2D class average, 3D classification and refinement are shown. (B) Gold standard FSC curves for the MT S-Trimer. (C) FSC curves of model-to-map.

**Fig. S8.**
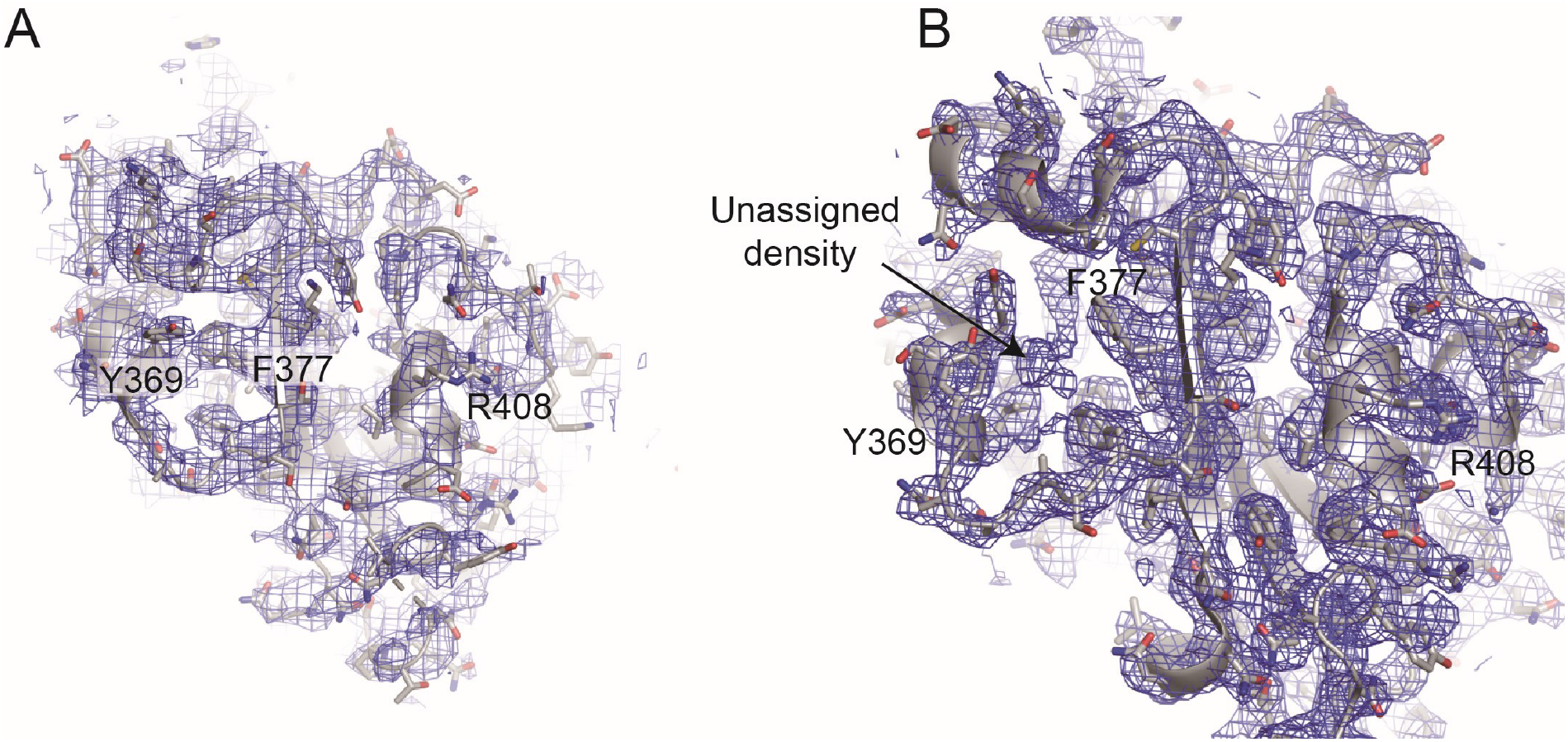
EM density map for RBD domains of the S-2P structure (PDB ID: 6VXX) (A) and the wildtype full-length structure (PDB ID: 6XR8).

**Table S1.**
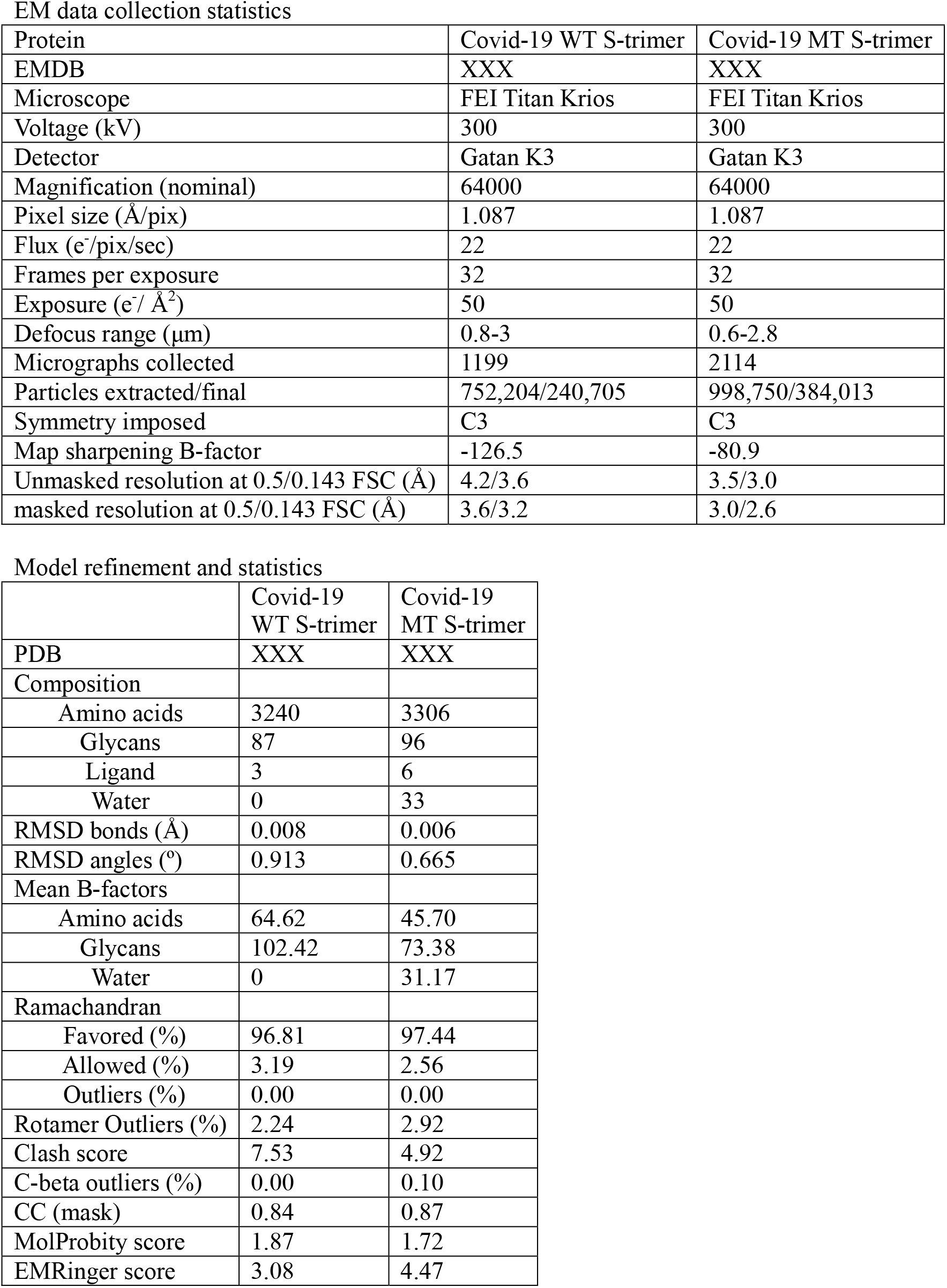
Cryo-EM data collection and refinement statistics.

EM data collection statistics
| Protein | Covid-19 WT S-trimer | Covid-19 MT S-trimer |
| --- | --- | --- |
| EMDB | XXX | XXX |
| Microscope | FEI Titan Krios | FEI Titan Krios |
| Voltage (kV) | 300 | 300 |
| Detector | Gatan K3 | Gatan K3 |
| Magnification (nominal) | 64000 | 64000 |
| Pixel size (Å/pix) | 1.087 | 1.087 |
| Flux (e<sup>-</sup>/pix/sec) | 22 | 22 |
| Frames per exposure | 32 | 32 |
| Exposure (e<sup>-</sup>/ Å<sup>2</sup>) | 50 | 50 |
| Defocus range (µm) | 0.8-3 | 0.6-2.8 |
| Micrographs collected | 1199 | 2114 |
| Particles extracted/final | 752,204/240,705 | 998,750/384,013 |
| Symmetry imposed | C3 | C3 |
| Map sharpening B-factor | -126.5 | -80.9 |
| Unmasked resolution at 0.5/0.143 FSC (Å) | 4.2/3.6 | 3.5/3.0 |
| masked resolution at 0.5/0.143 FSC (Å) | 3.6/3.2 | 3.0/2.6 |

Model refinement and statistics
|  | Covid-19<br>WT S-trimer | Covid-19<br>MT S-trimer |
| --- | --- | --- |
| PDB | XXX | XXX |
| Composition |  |  |
| Amino acids | 3240 | 3306 |
| Glycans | 87 | 96 |
| Ligand | 3 | 6 |
| Water | 0 | 33 |
| RMSD bonds (Å) | 0.008 | 0.006 |
| RMSD angles (°) | 0.913 | 0.665 |
| Mean B-factors |  |  |
| Amino acids | 64.62 | 45.70 |
| Glycans | 102.42 | 73.38 |
| Water | 0 | 31.17 |
| Ramachandran |  |  |
| Favored (%) | 96.81 | 97.44 |
| Allowed (%) | 3.19 | 2.56 |
| Outliers (%) | 0.00 | 0.00 |
| Rotamer Outliers (%) | 2.24 | 2.92 |
| Clash score | 7.53 | 4.92 |
| C-beta outliers (%) | 0.00 | 0.10 |
| CC (mask) | 0.84 | 0.87 |
| MolProbity score | 1.87 | 1.72 |
| EMRinger score | 3.08 | 4.47 |

**Table S2.**
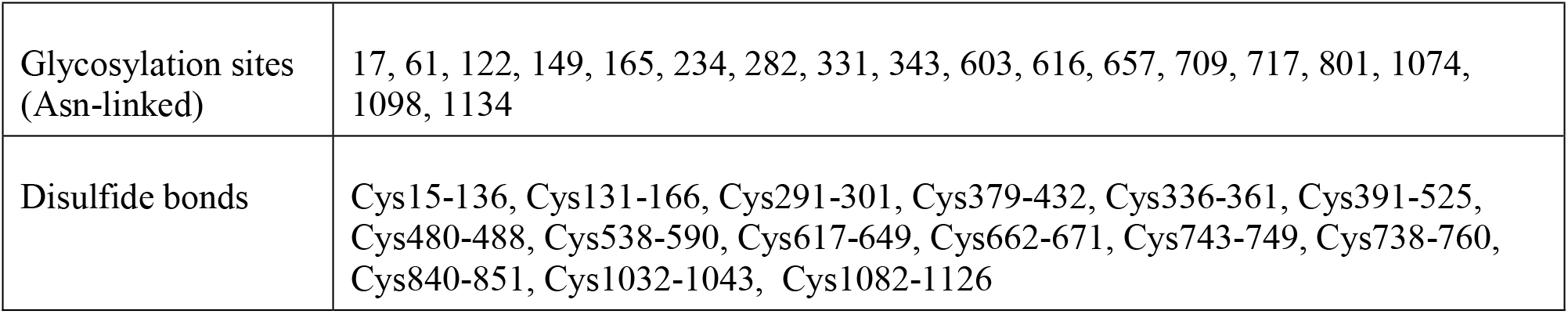
Summary of the glycosylation sites and disulfide bond observed in the structure.

|  |  |
| --- | --- |
| Glycosylation sites<br>(Asn-linked) | 17, 61, 122, 149, 165, 234, 282, 331, 343, 603, 616, 657, 709, 717, 801, 1074,<br>1098, 1134 |
| Disulfide bonds | Cys15-136, Cys131-166, Cys291-301, Cys379-432, Cys336-361, Cys391-525,<br>Cys480-488, Cys538-590, Cys617-649, Cys662-671, Cys743-749, Cys738-760,<br>Cys840-851, Cys1032-1043, Cys1082-1126 |

Movie S1. Conformational transition from the tightly closed state to open state

